# Complex regulation of Ca_v_2.2 N-type Ca^2+^ channels by Ca^2+^ and G-proteins Short title: Ca_v_2.2 modulation by Ca^2+^ and G-proteins

**DOI:** 10.1101/2024.11.19.624430

**Authors:** Jessica R. Thomas, Jinglang Sun, Juan De la Rosa Vazquez, Amy Lee

**Author notes:** Address correspondence to: Amy Lee, University of Texas-Austin Dept. of Neuroscience, 100 E. 24^th^ St, Austin, TX 78712.

## Abstract

G-protein coupled receptors inhibit Ca_v_2.2 N-type Ca^2+^ channels by a fast, voltage-dependent pathway mediated by Gα_i_/ Gβγ and a slow, voltage-independent pathway mediated by Gα_q_−dependent reductions in phosphatidylinositol 4,5-bisphosphate (PIP2) or increases in arachidonic acid. Studies of these forms of regulation generally employ Ba^2+^ as the permeant ion, despite that Ca^2+^ -dependent pathways may impinge upon G-protein modulation. To address this possibility, we compared tonic G-protein inhibition of currents carried by Ba^2+^ (*I_Ba_*) and Ca^2+^ (*I_Ca_*) in HEK293T cells transfected with Ca_v_2.2. Both *I_Ba_* and *I_Ca_* exhibited voltage-dependent facilitation (VDF), consistent with Gβγ unbinding from the channel. Compared to that for *I_Ba_*, VDF of *I_Ca_*was less sensitive to an inhibitor of Gα proteins (GDP-β-S) and an inhibitor of Gβγ (C-terminal construct of G-protein coupled receptor kinase 2). While insensitive to high intracellular Ca^2+^ buffering, VDF of *I_Ca_* that remained in GDP-β-S was blunted by reductions in PIP2. We propose that when G-proteins are inhibited, Ca^2+^ influx through Ca_v_2.2 promotes a form of VDF that involves PIP2. Our results highlight the complexity whereby Ca_v_2.2 channels integrate G-protein signaling pathways, which may enrich the information encoding potential of chemical synapses in the nervous system.

## INTRODUCTION

In nerve terminals, voltage-gated Ca_v_2 channels are prominent mediators of Ca^2+^ influx which triggers the exocytotic release of neurotransmitters into the synaptic cleft. The inhibition of presynaptic Ca_v_2 channels by neurochemicals such as GABA and norepinephrine potently suppresses neurotransmission via receptors coupled to heterotrimeric G-proteins (GPCRs) [1]. This inhibition can occur through a voltage-dependent, membrane-delimited pathway involving the Gα_i/o_ class of G-proteins and the binding of Gβγ to the channel [2–6]. GPCRs coupled to the Gα_q_ class of G-proteins also inhibit Ca_v_2 channels through a slower, voltage-independent pathway [7, 8]. Mechanisms for this form of Ca_v_2 channel modulation include enzymatic depletion of phosphatidylinositol 4,5-bisphosphate (PIP2) [9, 10], which normally enhances the activity of Ca_v_ channels [11].

Among the Ca_v_2 subtypes (Ca_v_2.1, Ca_v_2.2, Ca_v_2.3), Ca_v_2.2 channels exhibit particularly strong voltage-dependent inhibition by G-proteins [12, 13] which can be tempered by other signal mediators. For example, GPCRs that activate protein kinase C (PKC) diminish the impact of Gβγ on Ca_v_2.2 [14–16]. PKC phosphorylates a threonine in the cytoplasmic linker between domains I and II, which prevents the interaction with Gβγ [17, 18]. Conversely, several proteins involved with synaptic release, such as syntaxin 1A and cysteine string proteins, enhance G-protein inhibition of Ca_v_2.2 through interactions with both Gβγ and the channel [19–21]. Thus, the impact of GPCRs on neuronal Ca_v_2.2 channels may vary with patterns of neuronal activity, exposure to various neuromodulators, and interactions with proteins in specific subcellular compartments.

Ca_v_2.2 undergoes some voltage-dependent inhibition by G-proteins even without exogenous application of GPCR agonists, which could result from an excess of free Gβγ and/or activation of autoreceptor GPCRs [22–27]. In these studies, Ba^2+^ was often chosen as the permeant ion since Ba^2+^ currents (*I_Ba_*) are larger in amplitude than Ca^2+^ currents (*I_Ca_*) [28]. However, this approach can mask physiologically relevant forms of Ca_v_2.2 modulation that rely on Ca^2+^ influx [29] and could affect the impact of G-proteins. To address this possibility, we compared tonic G-protein modulation of *I_Ba_* and *I_Ca_* in HEK293T cells transfected with Ca_v_2.2. Our results indicate that tonic inhibition by Gβγ is stronger for *I_Ba_*than for *I_Ca_* and implicate PIP2 in modulation of *I_Ca_*and not *I_Ba_* when Gβγ-mediated inhibition is suppressed. Our findings add to the diverse modes by which Ca_v_ channels are regulated, some of which depend critically on the nature of the permeating cation.

## MATERIALS AND METHODS

*cDNAs.* The following cDNAs were used: Ca_v_2.2 e37b (Genbank # AF055477), β_2A_(Genbank # NM_053851), α_2_δ-1 (Genbank # NM_000722.3), pEGFP (Addgene). The C-terminal construct corresponding to G-protein coupled receptor kinase containing a myristic acid attachment signal (MAS-GRK2-ct) and zebrafish voltage-sensitive phosphatase (Dr-VSP) were described previously [8, 11].

### Cell culture and transfection

Human embryonic kidney 293 cells transformed with the SV40 T-antigen (HEK 293T, American Type Culture Collection Cat# CRL-3216, RRID:CVCL_0063) were maintained in Dulbecco’s modified Eagle’s medium with 10% fetal bovine serum and 1% penicillin-streptomycin at 37 °C in a humidified atmosphere with 5% CO_2_. Cells were grown to 80% confluence and transfected using Fugene 6 (Promega) according to the manufacturer’s protocol. Cells were plated in 35 mm dishes and transfected with cDNAs encoding Ca_v_ channel subunits (Ca_v_2.2, 1.8 μg; β_2A_, 0.6 μg; and α_2_δ-1, 0.6μg). In some experiments, 0.5 μg of MAS-GRK2-ct or Dr-VSP was co-transfected to buffer Gβγ or deplete PIP_2_, respectively. Cotransfection with cDNA encoding enhanced green fluorescent protein (pEGFP, 50 ng) allowed visualization of transfected cells.

### Electrophysiological recordings

Whole-cell patch recordings were performed 24-72 hours after transfection with a EPC-8patch clamp amplifier and Patch master software (HEKA Elektronik). External recoding solution contained (in mM): 150 Tris, 1 MgCl_2_, and 5 CaCl_2_ or BaCl_2_. Intracellular solution contained (in mM): 140 N-methyl-D-glucamine 10 HEPES, 10 EGTA, 2 MgCl_2_, and 2 Mg-ATP. The pH of both solutions was adjusted to 7.3 using methanesulfonic acid. In some experiments BAPTA or Guanosine5′-[β-thio]diphosphate trilithium salt (GDPβS) was added to the intracellular solution to either buffer Ca^2+^ or block G proteins, respectively. Electrode resistances were 4-6 MΩ in the bath solution. Series resistance was compensated 60-70%. Leak currents were subtracted using a P/-4 protocol. Data were analyzed using Igor Pro software (Wavemetrics). Averaged data represent mean ± S.E., and result from at least 3 independent transfections.

### Data presentation and statistical analysis

Data were incorporated into figures using Graph-Pad Prism software and Adobe Illustrator software. Statistical analysis was performed with Graph-Pad Prism software. The data were first analyzed for normality using the Shapiro–Wilk test. For parametric data, significant differences were determined by Student’s t test or ANOVA with post hoc Dunnett or Tukey test. For nonparametric data, the Mann-Whitney or Kruskal–Wallis with post hoc Dunn’s tests were used.

## RESULTS

In electrophysiological recordings of Ca_v_ channels, inhibition by G-proteins can be studied by evoking current-voltage (I-V) relationships before and after a depolarizing prepulse [23]. With this protocol, current amplitudes after the prepulse should be larger due to Gβγ unbinding from the channel [30]. We used this voltage protocol to test whether the tonic Ca_v_2.2 modulation by G-proteins might differ for *I_Ba_*and *I_Ca_* in transfected HEK293T cells. In our experiments, we used the Ca_v_2.2 splice variant containing exon 37b which lacks the voltage-independent, tyrosine kinase-dependent form of G-protein modulation seen for variants containing exon 37a [31]. We cotransfected cells with the auxiliary α_2_δ−1 subunit and β_2A_ subunit, which produces stronger tonic G-protein modulation than channels containing the β_1b_ subunit [23]. To account for differences in current amplitudes between cells due to variable levels of channel expression, we plotted I-V data normalized to the maximal current amplitude following the prepulse (*I_norm_*). As expected, the amplitudes of the normalized peak Ba^2+^ current (*I_norm_*, at test pulse = 0 mV) were significantly higher after (median = −0.84) than before a +60-mV prepulse (median = −0.33, W = −28, p = 0.02 by Wilcoxon matched-pairs test; Fig.1A-C). To verify the involvement of G-proteins, we used the guanosine diphosphate analog GDP-β-S which should limit the availability of Gβγ by stabilizing its association with Gα [32]. When GDP-β-S was included in the intracellular recording solution, the amplitude of peak *I_norm_* was still higher after (median = −0.58) than before the prepulse (median = −0.41, W = −28, p = 0.02 by Wilcoxon matched-pairs test; Fig.1D-F). However, the extent of the prepulse-induced increase in peak *I_norm_* was 10-fold lower with GDP-β-S (Fractional facilitation (FF) = 1.05 ± 0.27 for control vs. 0.2 ± 0.04 for +GDP-β-S, t = 3.081, df= 12, p = 0.01 by unpaired t-test; Fig.1G). These results show that Ca_v_2.2 undergoes tonic, voltage-dependent inhibition of Ca_v_2.2 by G-proteins in HEK293T cells, as described previously for this channel in other cell-types [23, 26].

**Fig. 1.**
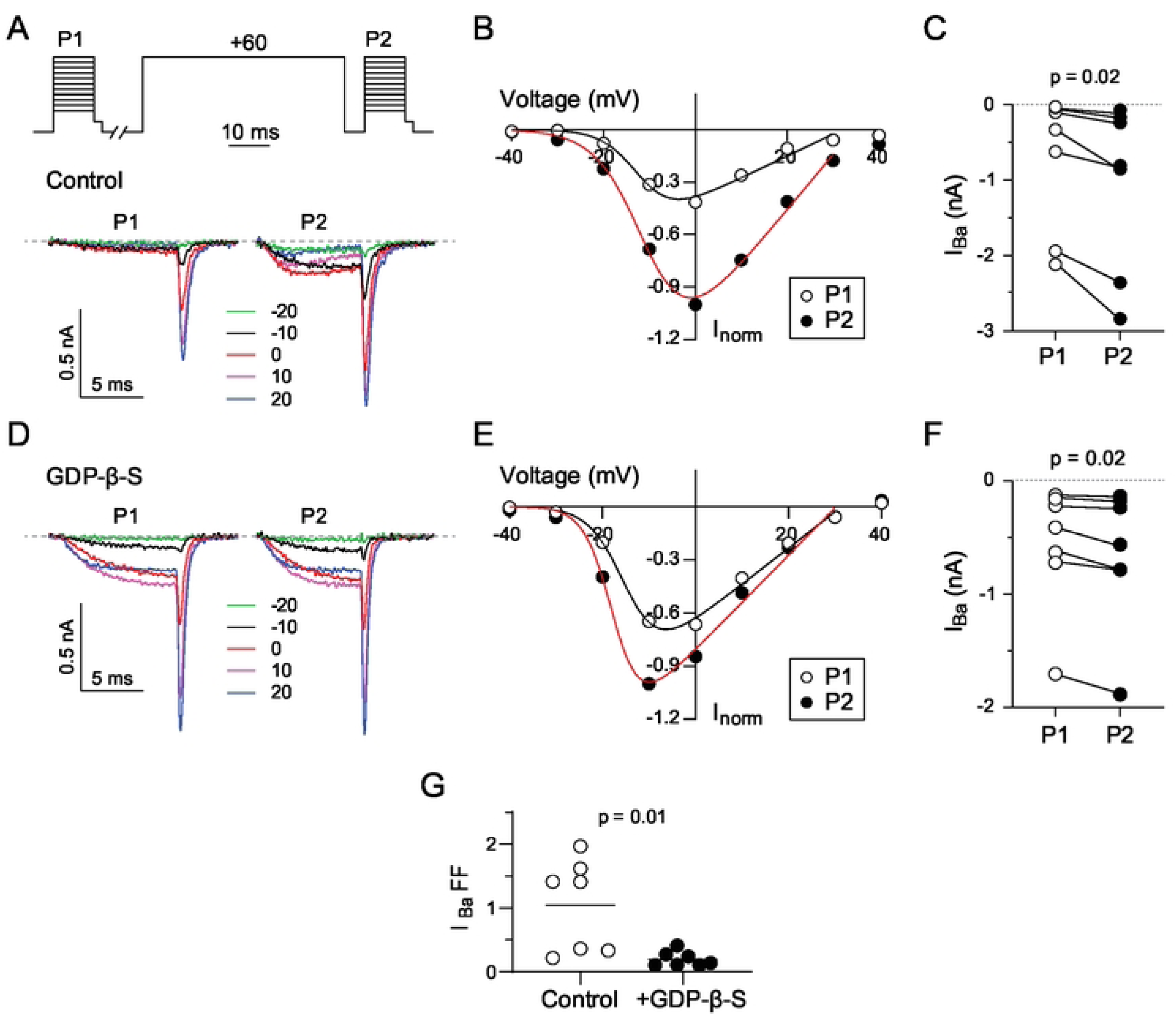
VDF of I_Ba_ for Ca_v_2.2 is blunted by GDPβS. (A) Representative current traces and voltage protocol. I_Ba_ was evoked by a 10-ms test pulse from a holding voltage of −80 mV to the indicated voltages 10 s before (P1) and 5 ms after (P2) a 50-ms conditioning pre-pulse to +60 mV. The test pulses were followed by a 2-ms step to −60 mV prior to repolarizing to −80 mV. (B) Representative I-V plot of P1 and P2 currents for a single cell. Smooth line represents Boltzmann fits. (C) I_Ba_ for P1 and P2 pulses for each cell. p-value was determined by Wilcoxon test and paired t-test. (D-F) Same as in A-C but for cells where GDPβS (0.3 mM) was included in the intracellular recording solution. (G) Plot comparing fractional facilitations, (P2-P1)/P1, for I_Ba_ evoked by 0 mV test pulse between cells with and without intracellular GDPβS. p-value was determined by unpaired t-test.

Like *I_Ba_, I_Ca_* also was increased by the prepulse under control conditions (median = −0.50 before vs. median = −0.74 after, t = 6.348, df = 6, p = 0.001; Fig.2A-C) and with GDP-β-S (median = −0.68 before vs. median = −0.82 after, W = −66, p = 0.001 by Wilcoxon matched-pairs test; Fig.2D-F). However, the depolarizing prepulse caused a significantly smaller increase in *I_Ca_* than *I_Ba_*under control conditions (FF = 0.35 ± 0.03 for *I_Ca_* vs. 1.05 ± 0.27 for *I_Ba_*, t = 2.549, df= 12, p = 0.02 by unpaired t-test). Moreover, facilitation caused by the prepulse did not significantly differ under control conditions and with GDP-β-S (FF = 0.25 ± 0.03, t = 2.007, df= 16, p = 0.06 compared to control by unpaired t-test; Fig.2G). Thus, tonic voltage-dependent inhibition of Ca_v_2.2 by G-proteins is weaker and less sensitive to GDP-β-S for *I_Ca_* than *I_Ba_*.

**Fig. 2.**
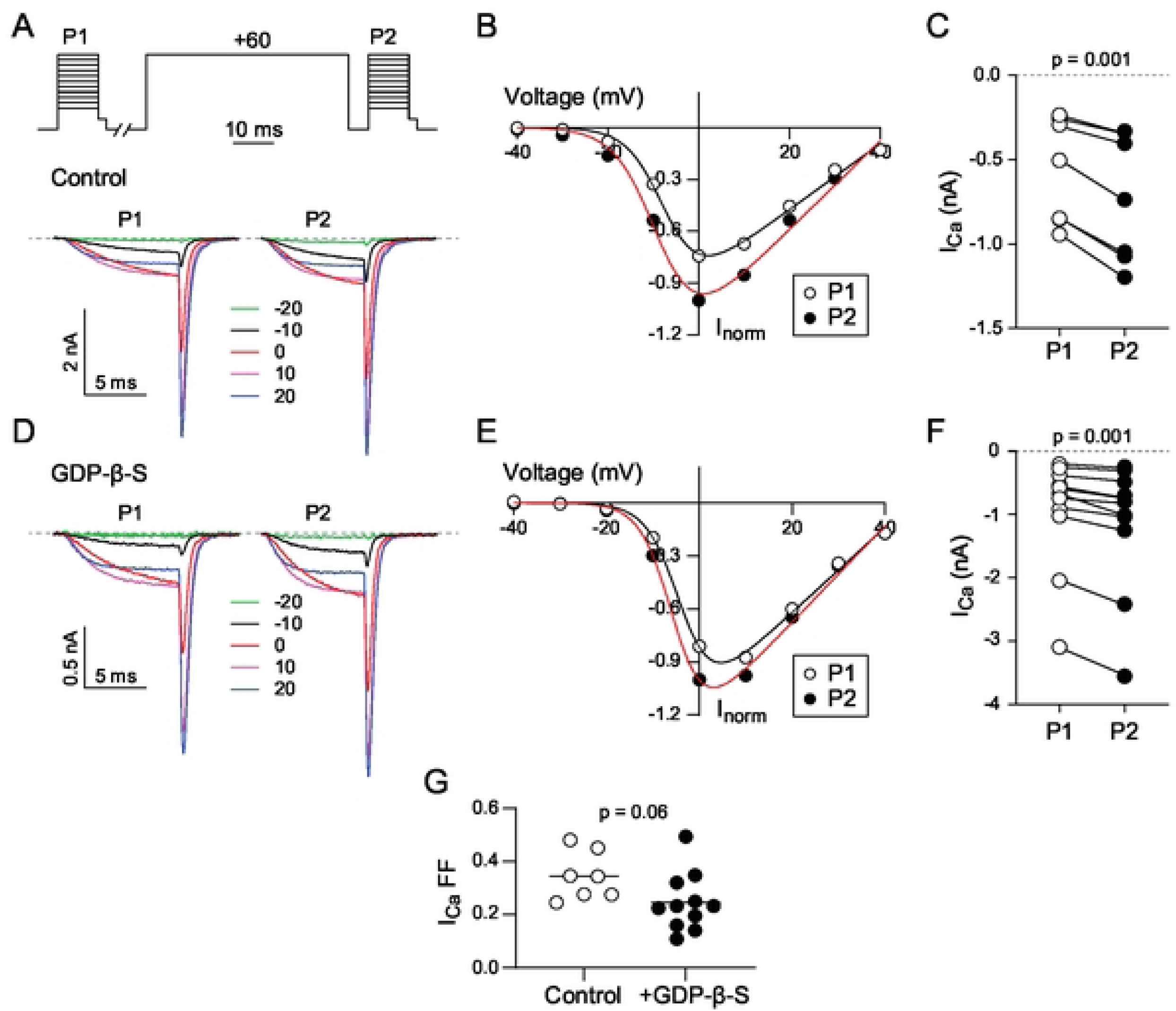
VDF of I_Ca_ for Ca_v_2.2 is unaffected by GDPβS. (A) Representative current traces and voltage protocol. I_Ca_ was evoked by the same voltage protocol as in Fig.1. (B) Representative I-V plot of P1 and P2 currents for a single cell. Smooth line represents Boltzmann fits. (C) I_Ca_ for P1 and P2 pulses for each cell. p-value was determined by paired t-test. (D-F) Same as in A-C but for cells where GDPβS (0.3 mM) was included in the intracellular recording solution. (G) Plot comparing fractional facilitations, (P2-P1)/P1, for I_Ca_ evoked by 0 mV test pulse between cells with and without intracellular GDPβS. p-value was determined by unpaired t-test.

To further investigate this difference in G-protein regulation of *I_Ca_* and *I_Ba_*, we used a double pulse protocol where the effect of varying the voltage of the prepulse is measured on a test current evoked before (P1) and after (P2) the prepulse (Fig.3A-F). With this protocol, VDF is evident as a progressive increase in the P2 vs P1 current amplitude with prepulse voltage [33]. For *I_Ba_*, VDF was robust under control conditions and was reduced by GDP-β-S (Fig.3B,D). The amount of VDF was measured as the difference in the P2 and P1 currents with a prepulse to +80 mV (Fractional facilitation, FF_80_) and was significantly lower with GDP-β-S (median FF_80_ = 0.14) compared to control conditions (median FF_80_ = 0.41, t = 3.748, df = 14, p = 0.0022 by unpaired t-test; Fig.3F). VDF of *I_Ca_* was also strong and showed a similar dependence on prepulse voltage as *I_Ba_.* However, unlike *I_Ba_,* VDF was not significantly different under control conditions (median FF_80_ = 0.52) and with GDP-β-S (median FF_80_ = 0.39, t = 2.076, df = 15, p = 0.056 by unpaired t-test; Fig.3C,E,F).

**Fig.3.**
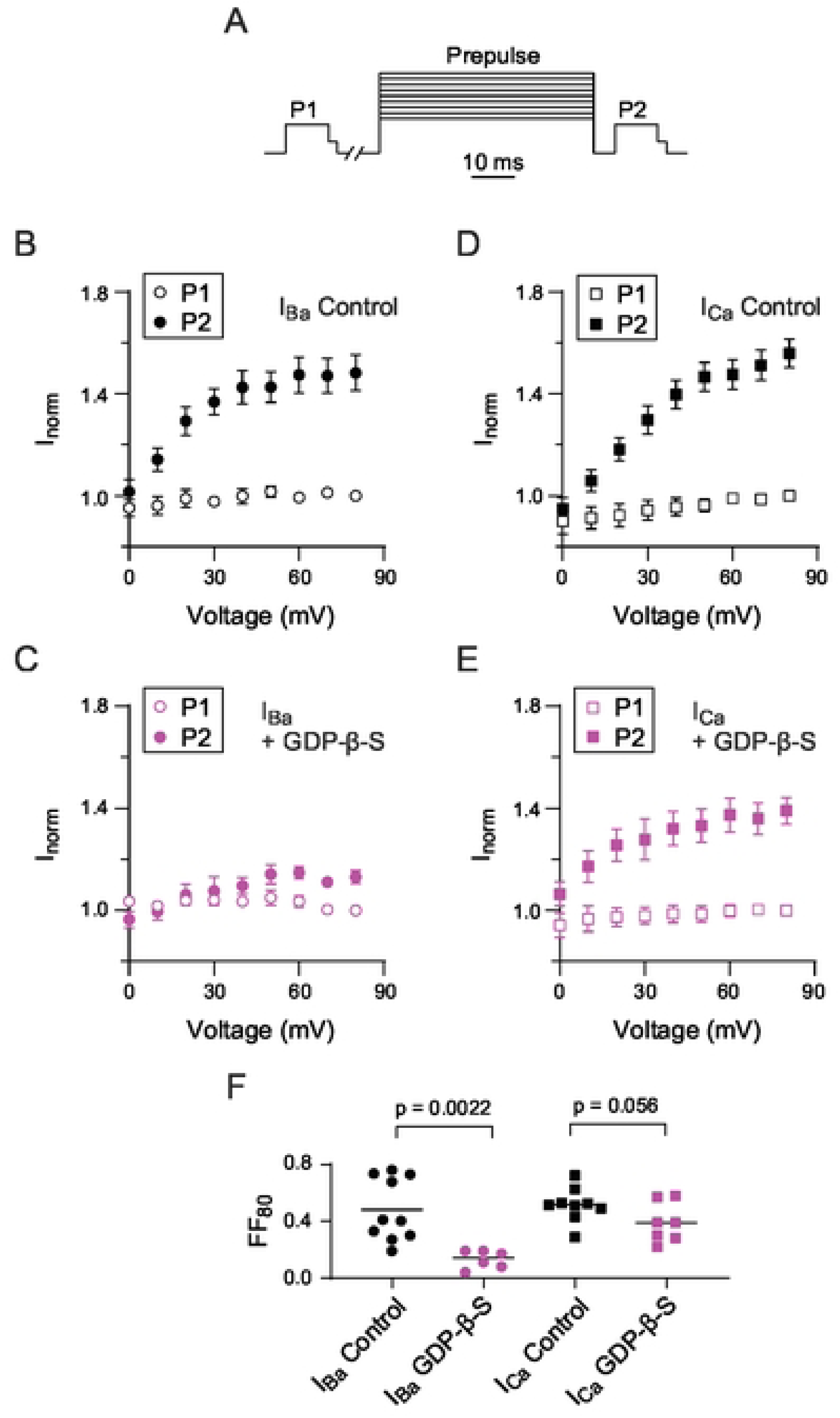
Decline of VDF of I_Ba_ for Ca_v_2.2 caused by GDPβS. (A) Voltage protocol. I_Ca_ (or I_Ba_) was evoked by a 10-ms test pulse from a holding voltage of −80 mV to −5 mV (−10 mV for I_Ba_) 10-s before (P1) and 5-ms after (P2) a 50-ms conditioning pre-pulse to indicated voltages. The test pulses were followed by a 2-ms step to −60 mV prior to repolarizing to −80 mV. (B-C) Normalized I_Ca_ and I_Ba_ evoked by P1 and P2 test pulse separated by variable conditioning pre pulses. The current value in each cell was normalized to current evoked by P1 test pulse before the conditioning pre-pulse at 80 mV. (D-E) Same as in B-C but for cells where GDPβS (0.3 mM) was included in the intracellular recording solution. (F) Plot comparing fractional facilitations, (P2-P1)/P1, for I_Ca_ and I_Ba_ evoked before and after a 80 mV conditioning pre pulse in cells with and without intracellular GDPβS. p-values were determined by the Mann-Whitney test.

A possible explanation for our results thus far was that VDF of *I_Ca_* could involve an additional pathway that is recruited even when Gβγ is inhibited. To test this, we coexpressed Ca_v_2.2 with a C-terminal construct of GPCR kinase 2 (GRK) which has no kinase activity but acts to sequester Gβγ [8]. With the I-V protocol to measure VDF, the peak current amplitude for both *I_Ca_* and *I_Ba_* was still increased by the +60 mV conditioning pulse in the presence of GRK (Fig.4A-F). As expected, VDF for *I_Ba_* was significantly weaker with GRK (FF = 0.19 ± 0.04, n = 7) than under control conditions (FF = 1.41 ± 0.27, n = 7; t = 3.102, df= 12, p = 0.009 by unpaired t-test; Fig.4G). In contrast, there was no significant difference in VDF for *I_Ca_* with GRK (FF = 0.32 ± 0.07, n = 5) than under control conditions (FF = 0.34 ± 0.03, n = 7; t = 0.314, df= 10, p = 0.759 by unpaired t-test; Fig.4G). Similar results were obtained with the double pulse protocol (Fig.5A-D). Compared to control conditions, GRK expression caused a significant reduction in VDF for *I_Ba_* (58%, Fig.5B,D) but a non-significant slight increase in VDF for *I_Ca_* (Fig.5C,D). These results agree with our hypothesis that VDF of *I_Ca_* could proceed even when Gβγ is inhibited.

**Fig. 4.**
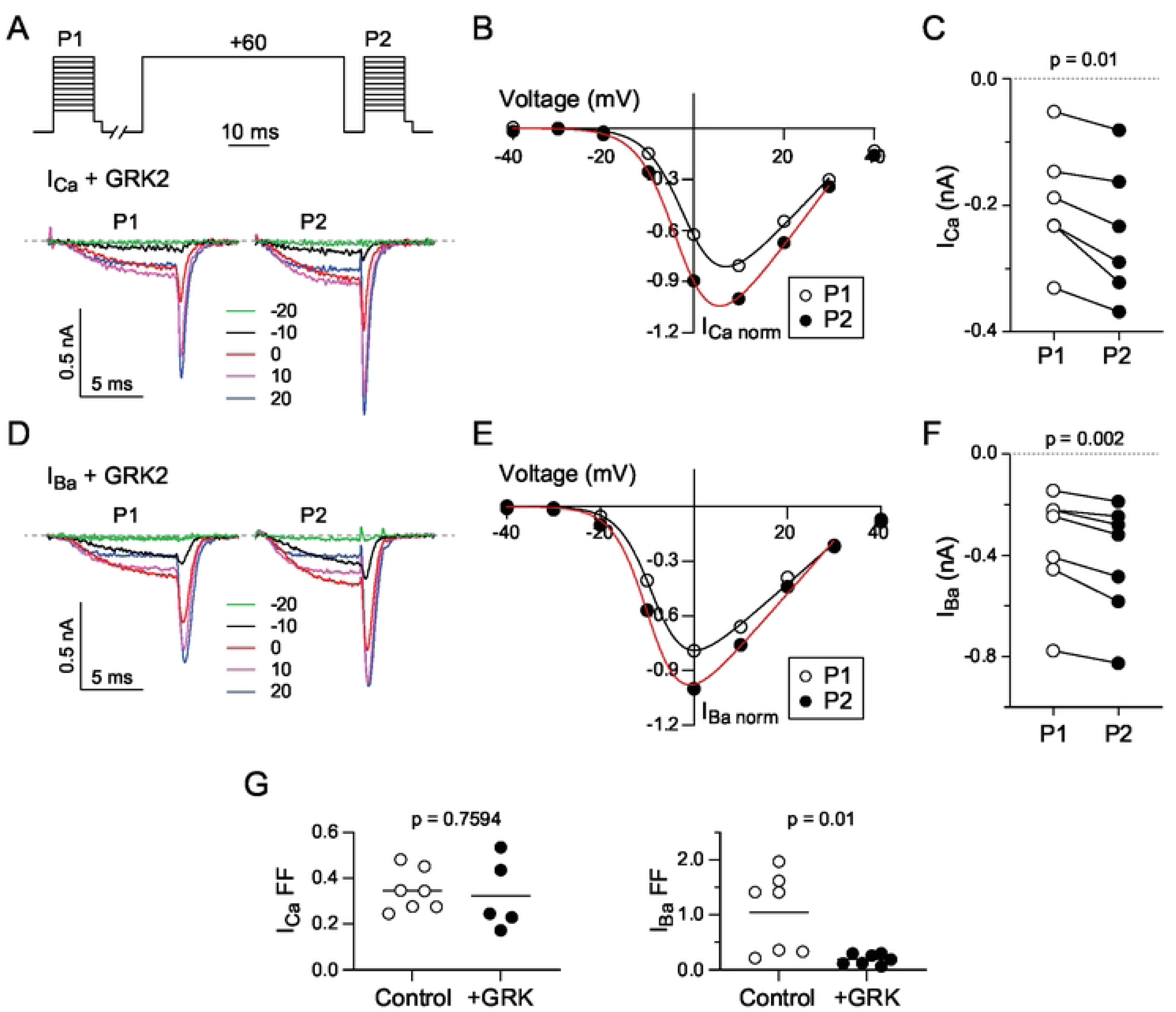
VDF of I_Ba_ for Ca_v_2.2 is suppressed by GRK2. (A) Representative current traces and voltage protocol. I_Ca_ was evoked by the same voltage protocol as in Fig.1. (B) Representative I-V plot of P1 and P2 currents for a single cell co-transfected with GRK2. Smooth line represents Boltzmann fits. (C) I_Ca_ for P1 and P2 pulses for each cell. p-value was determined by paired t-test. (D-F) Same as in A-C but for cells recorded in Ba²⁺ bath solution. (G) Plots comparing fractional facilitations, (P2-P1)/P1, for I_Ca_ and I_Ba_ evoked by 0 mV test pulse. P-values were determined by unpaired t-test.

**Figure 5.**
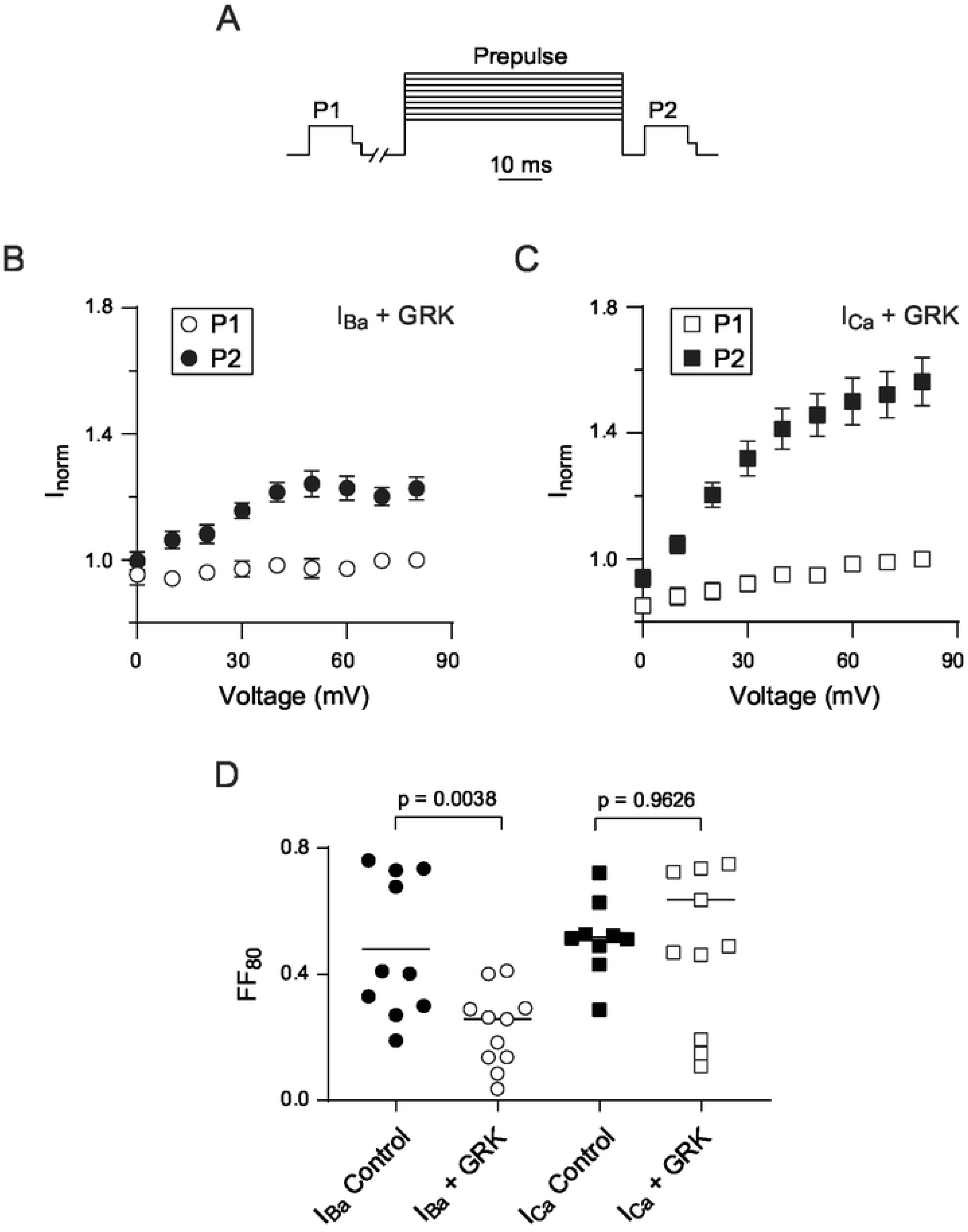
Reduced VDF of I_Ba_ for Ca_v_2.2 caused by GRK2. (A) Voltage protocol (same as in Fig.3). (B-C) Normalized I_Ca_ and I_Ba_ evoked by P1 and P2 test pulse separated by variable conditioning pre pulses. The current value in each cell was normalized to current evoked by P1 test pulse before the conditioning pre-pulse at 80 mV. Cells were co-transfected with GRK2. (D) Plot comparing fractional facilitations, (P2-P1)/P1, for I_Ca_ and I_Ba_ evoked before and after an 80 mV conditioning pre pulse in cells with and without GRK2 transfection. p-values were determined by unpaired t-test.

The apparent absence of an effect of GDPβS on VDF of *I_Ca_* (Figs.2,3) could signify opposing regulation of Ca_v_2.2 by another G-protein signaling pathway that is recruited when Ca^2+^ ions permeate the channel. One possibility was that GDP-β-S enabled a form of Ca^2+^-dependent facilitation (CDF) similar to that for Ca_v_2.1 channels that is mediated by calmodulin (CaM) binding to the Ca_v_2.1 C-terminal domain [34, 35]. This seemed unlikely since the VDF exhibited by Ca_v_2.2 *I_Ca_* in the double pulse protocol did not resemble CaM-dependent CDF of Ca_v_2.1, which shows a U-shaped dependence on prepulse voltage that reflects the amount of Ca^2+^ influx during the prepulse [36, 37]. Moreover, Ca_v_2.2 lacks key domains present in Ca_v_2.1 that are required for CaM-dependent CDF [38]. In primary sensory neurons, Ca_v_2.2 undergoes CDF that is mediated by CaM dependent protein kinase II (CaMKII), which requires cytoplasmic accumulation of Ca^2+^ [39]. In the voltage protocols for measuring VDF, the interval between the P1 and conditioning pulses is 10 s, which may allow for sufficient Ca^2+^ influx during the P1 pulse to activate Ca^2+^-dependent pathways such as those involving CaMKII. However, VDF of *I_Ca_* with strong Ca^2+^ buffering with 10 mM BAPTA (median FF_80_ = 0.38, Fig.6A,B) or the 0.3 mM of the CaMKII inhibitor KN93 (median FF_80_ = 0.32, Fig.6C) was similar to that with or without GDP-β-S (F(3, 17) = 1.781, p = 0.189 by One-Way ANOVA; Fig.6D). These results argue against a role for CaMKII in VDF of Ca_v_2.2 *I_Ca_*.

**Fig.6.**
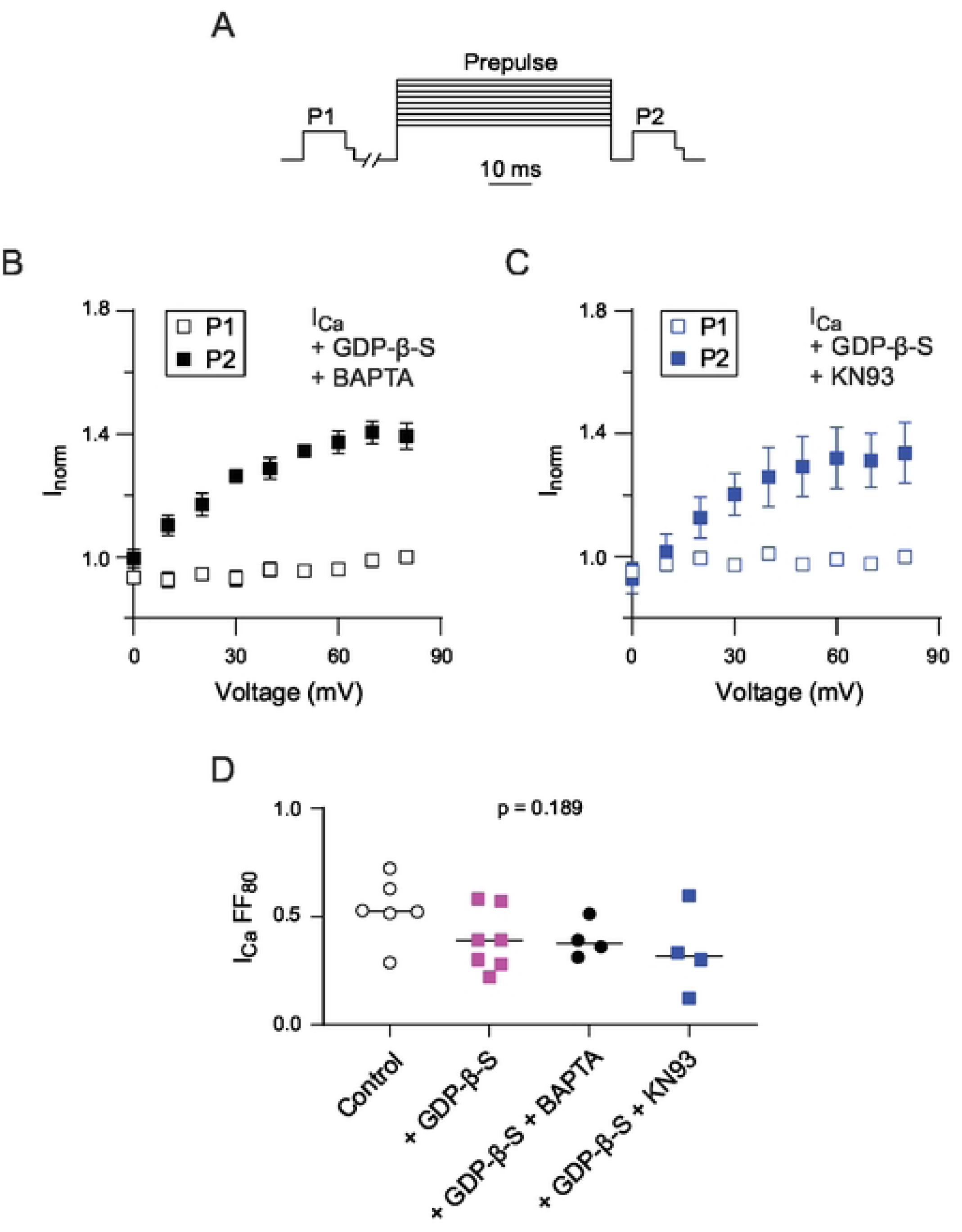
VDF of I_Ca_ for Ca_v_2.2 is not mediated by CaMKII. (A) Voltage protocol (same as in Fig.3). (B) Normalized I_Ca_ evoked by P1 and P2 test pulse separated by variable conditioning pre pulses. I_Norm_ was obtained by normalizing all I_Ca_ to I_Ca_ evoked by P1 test pulse before the conditioning pre-pulse at 80 mV. GDPβS (0.3 mM) and BAPTA (10 mM) was included in the intracellular recording solution. (C) Same as in B but for cells where GDPβS (0.3 mM) and KN93 was included in the intracellular recording solution. (D) Plot comparing fractional facilitations, (P2-P1)/P1, for I_Ca_ evoked before and after a 80 mV conditioning pre pulse in cells with various intracellular conditions. p-value was determined by One-Way ANOVA.

An alternative mechanism involves Gα_q_-dependent activation of phospholipase C (PLC), which causes the hydrolysis of PIP2 into inositol 1,4,5-trisphosphate and diacylglycerol, or increased liberation of arachidonic acid by phospholipase A2 [40]. It is well-established that PIP2 supports the function of Ca_v_ channels, and that GPCRs linked to Gα_q_ cause a decline in PIP2 that lowers activity of Ca_v_ channels [9, 11, 41, 42]. GDP-β-S might suppress this Gα_q_ signaling pathway, thus increasing Ca_v_ channel activity by reducing PIP2 hydrolysis. If selective for *I_Ca_*, this effect of GDP-β-S on Gα_q_ might mask the effect of GDP-β-S on the Gαi/ Gβγ-mediated pathway, leaving net VDF unchanged. To test this, we utilized a voltage-sensitive phosphatase (VSP) from zebrafish which enables the depletion of PIP2 in living cells following a strong depolarizing voltage step (i.e., +120 mV). This approach has been used previously to blunt Gα_q_-dependent inhibition of Ca_v_ channels [11].

To enable VSP activation, we modified our voltage protocol to include a +120-mV VSP-activating pulse prior to the P1 test pulse and VDF was measured as the ratio of the P2/P1 pulses with an intervening +20 mV prepulse (Fig.7A). A more modest depolarizing prepulse was used in these experiments to avoid additional activation of the VSP. GDPβS was included in the intracellular recording solution to replicate conditions that led to distinctions in VDF of *I_Ca_* and *I_Ba_* in Fig.3. When the double pulse protocol was given without the +120-mV pulse, P2/P1 for *I_Ca_* did not differ in cells with (median = 1.371) and without VSP (median = 1.471; t = 1.612, df = 12, p = 0.133 by unpaired t-test) indicating that VSP did not affect VDF when not activated. In cells transfected with VSP, P2/P1 for *I_Ca_* measured with the +120 mV pulse (median = 1.038) was significantly lower than when measured without the +120 mV pulse (median = 1.371; t = 8.264, df = 7, p < 0.0001 by paired t-test; Fig.7B,C). This result demonstrates that PIP2 enhances VDF of *I_Ca_*. As in control cells transfected with Ca_v_2.2 alone (i.e., -VSP), P2/P1 for *I_Ba_*(+VSP) was not significantly different with (median = 1.184) or without the +120 mV pulse (median = 1.344, W = −15, p = 0.426 by Wilcoxon matched pair signed rank test; Fig.7C,D,E). Therefore, alterations in PIP2 do not affect VDF for *I_Ba_*. Taken together, our results suggest that PIP2 enhances VDF of *I_Ca_* but not *I_Ba_*when G-proteins are inhibited.

**Fig. 7.**
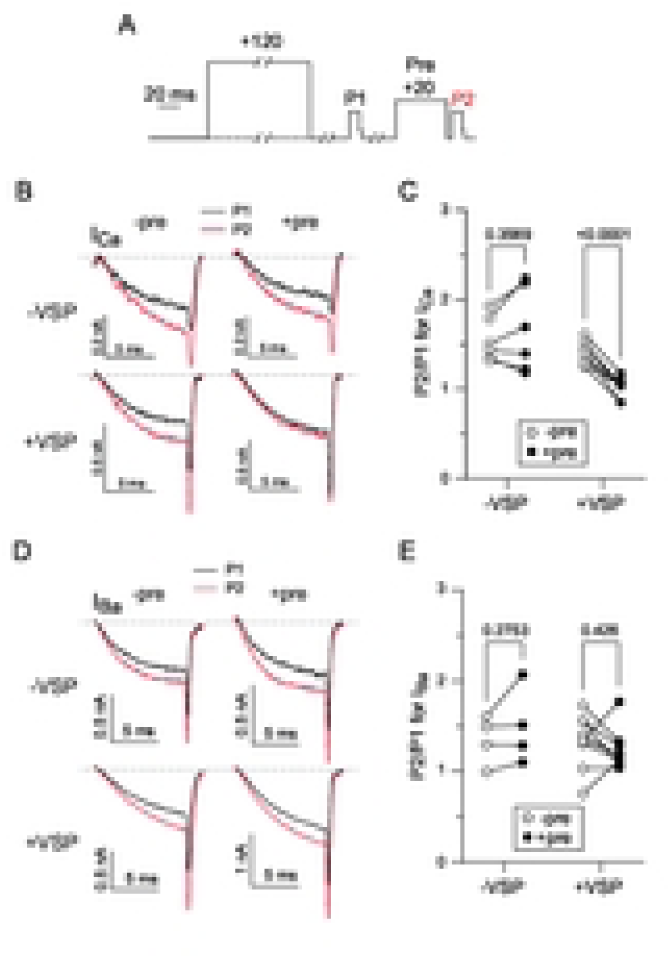
VDF of I_Ca_ for Ca_v_2.2 is abolished after PIP2 depletion via VSP activation. (A) Voltage protocol. An optional 1-s long +120 mV VSP activating pulse from a holding voltage of −80 mV is applied 10-s prior to the first test pulse. I_Ca_ or I_Ba_ was evoked by a 10-ms test pulse from a holding voltage of −80 mV to the indicated voltages (−5mV for I_Ca_ or −10mV for I_Ba_) 10-s before (P1) and 5-ms after (P2) a 50-ms conditioning pre-pulse to +20 mV. The test pulses were followed by a 2-ms step to −60 mV prior to repolarizing to −80 mV. (B) Paired representative I_Ca_ traces reflecting VDF from VSP and non-VSP transfected cells, with each cell tested with and without a +120 mV VSP activating pulse. (C) P2/P1 evoked I_Ca_ ratio for each cell. p-value was determined by paired t-test. (D) Same as in B-C but for cells recorded in Ba²⁺ bath solution. (E) P2/P1 evoked I_Ba_ ratio for each cell. p-value was determined by paired t-test (-VSP) and Wilcoxon test (+VSP).

## DISCUSSION

Our study reveals an unusual feature of G-protein modulation of Ca_v_2.2 that requires the influx of Ca^2+^ through the channel. For *I_Ba_*, VDF depends mainly on Gα_i_/ Gβγ which is blunted by GDPβS (Figs.1,3F, 8A) and GRK (Figs.4,5). For *I_Ca_*, VDF that remains in the presence of GDPβS (Figs.2,3) requires PIP2 since it is suppressed by Dr-VSP (Fig.7A-C). We propose that when Ca^2+^ permeates the channel, GDPβS inhibits not only Gα_i_/ Gβγ but also Gα_q_/ Gβγ. The latter pathway promotes a decline in PIP2 since both Gα_q_ and Gβγ can activate PLC [43]. Despite the competing effects of blunting the Gα_i_/ Gβγ pathway, GDPβS strengthens VDF of *I_Ca_*by limiting Gα_q_-dependent reductions in PIP2 (Fig.8B).

**Fig. 8.**
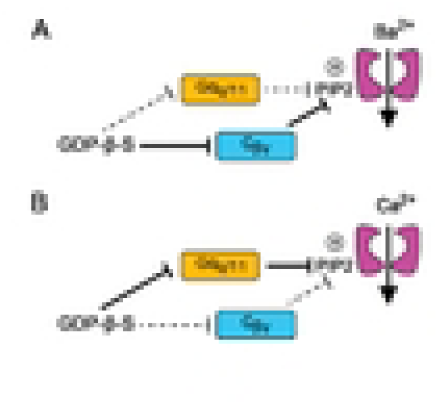
Model for distinct G-protein modulation of *I_Ba_* and *I_Ca_* mediated by Ca_v_2.2. (A,B) Ca_v_2.2 channels are potentiated by PIP2 binding and inhibited by Gβγ binding to the channel. Gα_q_-mediated decreases in PIP2 would be expected to inhibit *I_Ba_* and *I_Ca_*, whereas Gβγ would be expected to promote VDF. For *I_Ba,_* the main effect of GDP-β-S is to suppress VDF by inhibiting liberation of Gβγ from Gα_i_ (A). For *I_Ca_,* the main effect of GDP-β-S is to increase VDF by inhibiting Gα_q_-mediated decreases in PIP2 (B).

In studies of Ca_v_ channels, Ba^2+^ is often substituted for Ca^2+^ in the extracellular recording solution in part to minimize Ca^2+^-dependent pathways that could complicate analysis of intrinsic channel properties. However, *I_Ca_* can differ from *I_Ba_* in physiologically relevant ways. A prominent example is Ca^2+^-dependent inactivation (CDI), which is a characteristic of all Ca_v_1 and Ca_v_2 channels and manifests as faster decay of *I_Ca_* compared to *I_Ba_* [37, 44]. Ca_v_2 channels also undergo CDF [34–36], which for Ca_v_2.2 requires CaMKII and is reduced following nerve injury [39]. The VDF of *I_Ca_* for Ca_v_2.2 in our study differed from CaMKII-dependent CDF since it was not blocked by high BAPTA or the CaMKII inhibitor (Fig.6). The BAPTA-insensitivity suggests that Ca^2+^ elevations within a nanodomain of the Ca_v_2.2 channel are needed to support VDF when G-proteins are inhibited. Considering that Ca^2+^ can increase the enzymatic activity of PLC [45, 46], Ca^2+^ influx through Ca_v_2.2 could amplify the effects of Gα_q_ coupling to PLC, leading to greater reductions in PIP2 levels and reduced channel function than when using Ba^2+^ as the permeant ion. The persistence of VDF in the presence of GDPβS could then be viewed as a disinhibition of Ca_v_2.2 by stabilizing PIP2 levels that support channel function (Fig.8B). Some PLC isoforms are membrane-associated and can form macromolecular complexes with ion channels and GPCRs to allow for fast and localized signaling [47, 48]. Ca_v_ channels interact with a variety of proteins [1] including those that may scaffold PLC and position it for regulation by incoming Ca^2+^ ions. In addition, micromolar concentrations of Ca^2+^ can cluster PIP2 in nanodomains [49, 50] which might make PIP2 a more appealing substrate for hydrolysis by PLC than in the presence of Ba^2+^ ions.

PIP2 binds to a site in domain II S4 [51, 52] of Ca_v_2.2 which is critical for the ability of PIP2 to support channel function [53]. PIP2 also binds to a site in the cytoplasmic I-II linker of Ca_v_2.2, which is important for PIP2 modulation of Ca_v_2.2 channels containing the auxiliary β_2c_ but not β_2a_ subunit [53]. For this reason, Ca_v_2.2 channels containing β_2a_, such as those in our study, are less sensitive to PIP2 modulation, which may explain our findings that *I_Ba_* was not as significantly affected by Dr-VSP activation as shown previously [11, 53]. Studies of PIP2 modulation of Ca_v_2 channels have mainly focused on how its depletion promotes a decline in current density and increases in inactivation. However, PIP2 has been shown to support VDF of Ca_v_2.2 channels in hypothalamic neurons [54]. Future studies are needed to dissect the mechanisms whereby PIP2 binding enables gating transitions that underlie VDF when G-protein-mediated inhibition is suppressed.

## REFERENCES

1. Dolphin AC, Lee A. Presynaptic calcium channels: specialized control of synaptic neurotransmitter release. Nat Rev Neurosci. 2020. doi: 10.1038/s41583-020-0278-2.

2. Herlitze S, Garcia DE, Mackie K, Hille B, Scheuer T, Catterall WA. Modulation of Ca2+ channels by G-protein beta gamma subunits. Nature. 1996;380(6571):258-62. doi: 10.1038/380258a0.

3. Ikeda SR. Voltage-dependent modulation of N-type calcium channels by G-protein beta gamma subunits. Nature. 1996;380(6571):255-8. doi: 10.1038/380255a0.

4. Agler HL, Evans J, Tay LH, Anderson MJ, Colecraft HM, Yue DT. G protein-gated inhibitory module of N-type Ca_v_2.2 Ca^2+^ channels. Neuron. 2005;46(6):891–904.

5. Canti C, Bogdanov Y, Dolphin AC. Interaction between G proteins and accessory subunits in the regulation of 1B calcium channels in Xenopus oocytes. J Physiol. 2000;527 Pt 3:419–32.

6. Zamponi GW, Snutch TP. Decay of prepulse facilitation of N type calcium channels during G protein inhibition is consistent with binding of a single Gbeta subunit. Proc Natl Acad Sci U S A. 1998;95(7):4035–9.

7. Liu L, Zhao R, Bai Y, Stanish LF, Evans JE, Sanderson MJ, et al. M1 muscarinic receptors inhibit L-type Ca^2+^ current and M-current by divergent signal transduction cascades. J Neurosci. 2006;26(45):11588–98.

8. Kammermeier PJ, Ikeda SR. Expression of RGS2 alters the coupling of metabotropic glutamate receptor 1a to M-type K^+^ and N-type Ca^2+^ channels. Neuron. 1999;22:819–29.

9. Gamper N, Reznikov V, Yamada Y, Yang J, Shapiro MS. Phosphatidylinositol [correction] 4,5-bisphosphate signals underlie receptor-specific Gq/11-mediated modulation of N-type Ca2+ channels. J Neurosci. 2004;24(48):10980–92. doi: 10.1523/JNEUROSCI.3869-04.2004.

10. Liu L, Bonventre JV, Rittenhouse AR. cPLA2alpha-/-sympathetic neurons exhibit increased membrane excitability and loss of N-Type Ca2+ current inhibition by M1 muscarinic receptor signaling. PLoS One. 2018;13(12):e0201322. doi: 10.1371/journal.pone.0201322.

11. Suh BC, Leal K, Hille B. Modulation of high-voltage activated Ca^2+^ channels by membrane phosphatidylinositol 4,5-bisphosphate. Neuron. 2010;67(2):224–38. doi: 10.1016/j.neuron.2010.07.001.

12. Currie KP, Fox AP. Comparison of N- and P/Q-type voltage-gated calcium channel current inhibition. J Neurosci. 1997;17(12):4570–9.

13. Colecraft HM, Patil PG, Yue DT. Differential occurrence of reluctant openings in G-protein-inhibited N- and P/Q-type calcium channels. J Gen Physiol. 2000;115(2):175–92.

14. Swartz KJ, Merritt A, Bean BP, Lovinger DM. Protein kinase C modulates glutamate receptor inhibition of Ca^2+^ channels and synaptic transmission. Nature. 1993;361:165–8.

15. Zhu Y, Ikeda SR. VIP inhibits N-type Ca^2+^ channels of symnapthetic neurons via a pertussis toxin-insensitive but cholera toxin-sensitive pathway. Neuron. 1994;13:657–69.

16. Barrett CF, Rittenhouse AR. Modulation of N-type calcium channel activity by G-proteins and protein kinase C. J Gen Physiol. 2000;115(3):277–86. doi: 10.1085/jgp.115.3.277.

17. Zamponi GW, Bourinet E, Nelson D, Nargeot J, Snutch TP. Crosstalk between G proteins and protein kinase C mediated by the calcium channel α_1_ subunit. Nature. 1997;385:442–6.

18. Cooper CB, Arnot MI, Feng ZP, Jarvis SE, Hamid J, Zamponi GW. Cross-talk between G-protein and protein kinase C modulation of N-type calcium channels is dependent on the G-protein beta subunit isoform. J Biol Chem. 2000;275(52):40777–81. doi: 10.1074/jbc.C000673200.

19. Lu Q, AtKisson MS, Jarvis SE, Feng ZP, Zamponi GW, Dunlap K. Syntaxin 1A supports voltage-dependent inhibition of alpha1B Ca^2+^ channels by Gβ*γ* in chick sensory neurons. J Neurosci. 2001;21(9):2949–57.

20. Jarvis SE, Zamponi GW. Distinct molecular determinants govern syntaxin 1A-mediated inactivation and G-protein inhibition of N-type calcium channels. J Neurosci. 2001;21(9):2939–48.

21. Magga JM, Jarvis SE, Arnot MI, Zamponi GW, Braun JE. Cysteine string protein regulates G protein modulation of N-type calcium channels. Neuron. 2000;28(1):195–204.

22. Doupnik CA, Pun RY. G-protein activation mediates prepulse facilitation of Ca^2+^ channel currents in bovine chromaffin cells. J Membr Biol. 1994;140(1):47–56. doi: 10.1007/BF00234485.

23. Stephens GJ, Brice NL, Berrow NS, Dolphin AC. Facilitation of rabbit alpha1B calcium channels: involvement of endogenous Gβγ subunits. J Physiol. 1998;509 ( Pt 1):15–27.

24. Li B, Zhong H, Scheuer T, Catterall WA. Functional role of a C-terminal Gβγ-binding domain of Ca_v_2.2 channels. Mol Pharmacol. 2004;66(3):761–9. doi: 10.1124/mol.66.3.

25. Zhong H, Li B, Scheuer T, Catterall WA. Control of gating mode by a single amino acid residue in transmembrane segment IS3 of the N-type Ca^2+^ channel. Proc Natl Acad Sci U S A. 2001;98(8):4705–9.

26. Kasai H. Tonic inhibition and rebound facilitation of a neuronal calcium channel by a GTP-binding protein. Proc Natl Acad Sci USA. 1991;88:8855–9.

27. Quiroz-Acosta T, Bermeo K, Arenas I, Garcia DE. G-protein tonic inhibition of calcium channels in pancreatic beta-cells. Am J Physiol Cell Physiol. 2023;325(3):C592–C8. doi: 10.1152/ajpcell.00447.2022.

28. McNaughton NC, Randall AD. Electrophysiological properties of the human N-type Ca^2+^ channel: I. Channel gating in Ca^2+^, Ba^2+^ and Sr^2+^ containing solutions. Neuropharmacology. 1997;36(7):895–915. doi: 10.1016/s0028-3908(97)00085-3.

29. Christel C, Lee A. Ca^2+^-dependent modulation of voltage-gated Ca^2+^ channels. Biochim Biophys Acta. 2012;1820:1243–52. doi: 10.1016/j.bbagen.2011.12.012.

30. Zamponi GW, Snutch TP. Decay of prepulse facilitation of N type calcium channels during G protein inhibition is consistent with binding of a single Gβγ subunit. Proc Natl Acad Sci USA. 1998;95:4035–9.

31. Raingo J, Castiglioni AJ, Lipscombe D. Alternative splicing controls G protein-dependent inhibition of N-type calcium channels in nociceptors. Nat Neurosci. 2007;10(3):285–92. doi: 10.1038/nn1848.

32. Eckstein F, Cassel D, Levkovitz H, Lowe M, Selinger Z. Guanosine 5’-O-(2-thiodiphosphate). An inhibitor of adenylate cyclase stimulation by guanine nucleotides and fluoride ions. J Biol Chem. 1979;254(19):9829–34.

33. Ikeda SR. Double-pulse calcium channel current facilitation in adult rat sympathetic neurones. JPhysiol. 1991;439:181–214.

34. Lee A, Wong ST, Gallagher D, Li B, Storm DR, Scheuer T, et al. Ca^2+^/calmodulin binds to and modulates P/Q-type calcium channels. Nature. 1999;399(6732):155-9.

35. DeMaria CD, Soong T, Alseikhan BA, Alvania RS, Yue DT. Calmodulin bifurcates the local Ca^2+^ signal that modulates P/Q-type Ca^2+^ channels. Nature. 2001;411:484–9.

36. Lee A, Scheuer T, Catterall WA. Ca^2+^/calmodulin-dependent facilitation and inactivation of P/Q-type Ca^2+^ channels. J Neurosci. 2000;20(18):6830–8.

37. Thomas JR, Lee A. Measuring Ca^2+^-Dependent Modulation of Voltage-Gated Ca^2+^ Channels in HEK-293T Cells. Cold Spring Harb Protoc. 2016;2016(9):pdb prot087213. doi: 10.1101/pdb.prot087213.

38. Thomas JR, Hagen J, Soh D, Lee A. Molecular moieties masking Ca(2+)-dependent facilitation of voltage-gated Cav2.2 Ca(2+) channels. J Gen Physiol. 2018;150(1):83–94. doi: 10.1085/jgp.201711841.

39. Tang Q, Bangaru ML, Kostic S, Pan B, Wu HE, Koopmeiners AS, et al. Ca^2+^-dependent regulation of Ca^2+^ currents in rat primary afferent neurons: role of CaMKII and the effect of injury. J Neurosci. 2012;32(34):11737–49. doi: 10.1523/JNEUROSCI.0983-12.2012.

40. Roberts-Crowley ML, Mitra-Ganguli T, Liu L, Rittenhouse AR. Regulation of voltage-gated Ca2+ channels by lipids. Cell calcium. 2009;45(6):589–601. doi: 10.1016/j.ceca.2009.03.015.

41. Wu L, Bauer CS, Zhen XG, Xie C, Yang J. Dual regulation of voltage-gated calcium channels by PtdIns(4,5)P2. Nature. 2002;419(6910):947-52. doi: 10.1038/nature01118.

42. Vivas O, Castro H, Arenas I, Elias-Vinas D, Garcia DE. PIP(2) hydrolysis is responsible for voltage independent inhibition of CaV2.2 channels in sympathetic neurons. Biochemical and biophysical research communications. 2013;432(2):275–80. doi: 10.1016/j.bbrc.2013.01.117.

43. Pfeil EM, Brands J, Merten N, Vogtle T, Vescovo M, Rick U, et al. Heterotrimeric G Protein Subunit Galphaq Is a Master Switch for Gbetagamma-Mediated Calcium Mobilization by Gi-Coupled GPCRs. Molecular cell. 2020;80(6):940–54 e6. doi: 10.1016/j.molcel.2020.10.027.

44. Liang H, DeMaria CD, Erickson MG, Mori MX, Alseikhan B, Yue DT. Unified mechanisms of Ca^2+^ regulation across the Ca^2+^ channel family. Neuron. 2003;39:951–60.

45. Horowitz LF, Hirdes W, Suh BC, Hilgemann DW, Mackie K, Hille B. Phospholipase C in living cells: activation, inhibition, Ca2+ requirement, and regulation of M current. J Gen Physiol. 2005;126(3):243–62. doi: 10.1085/jgp.200509309.

46. Hashimotodani Y, Ohno-Shosaku T, Tsubokawa H, Ogata H, Emoto K, Maejima T, et al. Phospholipase Cbeta serves as a coincidence detector through its Ca2+ dependency for triggering retrograde endocannabinoid signal. Neuron. 2005;45(2):257–68. doi: 10.1016/j.neuron.2005.01.004.

47. Jiang Y, Wang S, Holcomb J, Trescott L, Guan X, Hou Y, et al. Crystallographic analysis of NHERF1-PLCbeta3 interaction provides structural basis for CXCR2 signaling in pancreatic cancer. Biochemical and biophysical research communications. 2014;446(2):638–43. doi: 10.1016/j.bbrc.2014.03.028.

48. Suh PG, Hwang JI, Ryu SH, Donowitz M, Kim JH. The roles of PDZ-containing proteins in PLC-beta-mediated signaling. Biochemical and biophysical research communications. 2001;288(1):1–7. doi: 10.1006/bbrc.2001.5710.

49. Han K, Kim SH, Venable RM, Pastor RW. Design principles of PI(4,5)P(2) clustering under protein-free conditions: Specific cation effects and calcium-potassium synergy. Proc Natl Acad Sci U S A. 2022;119(22):e2202647119. doi: 10.1073/pnas.2202647119.

50. Wen Y, Vogt VM, Feigenson GW. Multivalent Cation-Bridged PI(4,5)P(2) Clusters Form at Very Low Concentrations. Biophys J. 2018;114(11):2630–9. doi: 10.1016/j.bpj.2018.04.048.

51. Dong Y, Gao Y, Xu S, Wang Y, Yu Z, Li Y, et al. Closed-state inactivation and pore-blocker modulation mechanisms of human Ca(V)2.2. Cell rep. 2021;37(5):109931. doi: 10.1016/j.celrep.2021.109931.

52. Gao S, Yao X, Yan N. Structure of human Ca(v)2.2 channel blocked by the painkiller ziconotide. Nature. 2021;596(7870):143-7. doi: 10.1038/s41586-021-03699-6.

53. Park CG, Yu W, Suh BC. Molecular basis of the PIP(2)-dependent regulation of Ca(V)2.2 channel and its modulation by Ca(V) beta subunits. Elife. 2022;11. doi: 10.7554/eLife.69500.

54. Kirchner MK, Armstrong WE, Guan D, Ueta Y, Foehring RC. PIP(2) alters of Ca(2+) currents in acutely dissociated supraoptic oxytocin neurons. Physiol Rep. 2019;7(16):e14198. doi: 10.14814/phy2.14198.

